# SonoNERFs: Neural Radiance Fields applied to Biological Echolocation Systems allow 3D Scene Reconstruction Through Perceptual Prediction

**DOI:** 10.1101/2024.04.20.590416

**Authors:** Wouter Jansen, Jan Steckel

## Abstract

In this paper, we introduce SonoNERFs, a novel approach that adapts Neural Radiance Fields (NeRFs) to model and understand the echolocation process in bats, focusing on the challenges posed by acoustic data interpretation without phase information. Leveraging insights from the field of optical NeRFs, our model, termed SonoNERF, represents the acoustic environment through Neural Reflectivity Fields. This model allows us to reconstruct three-dimensional scenes from echolocation data, obtained by simulating how bats perceive their surroundings through sound. By integrating concepts from biological echolocation and modern computational models, we demonstrate the SonoNERF’s ability to predict echo spectrograms for unseen echolocation poses and effectively reconstruct a mesh-based and energy-based representation of complex scenes. Our work bridges a gap in understanding biological echolocation and proposes a methodological framework that provides a first order model on how scene understanding might arise in echolocating animals. We demonstrate the efficacy of the SonoNERF model on three scenes of increasing complexity, including some biologically relevant prey-predator interactions.

## I. Introduction

Echolocating bats exhibit a strong ability to localize and discriminate prey objects using sound as the primary sensing modality. A subset of bats called gleaning bats, are especially adept at finding and identifying prey in dense clutter [1], [2], [3], [4], [5], [6]. One of the main theories that explain this behavior is that these animals make clever use of physical interactions of their echolocation signals and the clutter in which their prey is perched upon [7], [8], [9], [10], [11]. Another class of bats called nectar-feeding bats, locate the flowers from which they nourish themselves using a special kind of leaf that is co-evolved by the pitcher plants that bear these flowers [12], [13], [14]. Similar traits of co-evolution have also been observed in other plant-bat relationships, such as bat-pollinated cacti [15]. In these plant-bat interactions, physical interactions between the emitted sound signals and the objects of interest give rise to emergent, stable spectral cues, which the bat can utilize to localize these leaves [14]. These insights have led to robotic applications, building a specific set of sonar reflectors that can be localized reliably by manufactured sonar sensors [16], [17].

While there is a growing body of evidence of the prevalence of these robust emergent cues, which bats utilize to solve their prey localization and identification task, the hypothesis of more in-depth scene understanding and modeling in bats still lingers. Indeed, when visiting bat research conferences, talks about 3D scene models in bats can often be overheard, giving rise to intense discussions about the sense or nonsense of such models existing. This is understandable, as humans like to reason in high-level 3D models of the environment, as this representation is natural to us. However, it is essential to understand that there are significant differences between the sensory data originating from a complex scene when sensed using optical sensors (with wavelengths in the range of hundreds of nanometers) or with acoustic sensors (using wavelengths in the order of several millimeters [18]).

This prevalence of the concept of an “acoustic image” or “acoustic 3D model” is not unsurprising, given the vast amount of literature by the pioneering researchers of bat echolocation [19], [20], [21], [22], [23]. Furthermore, some researchers have proposed systems that perform tomographic reconstruction of complex scenes using echolocation-like signals and use these generated images to explain certain phenomena observed in bat-prey interactions [24], [25], [26]. Many of these previous works consider the problem of image formation based on the reception time-domain reception signal, which represents the acoustic wave field impinging on the external ears of the bats. However, an important note here is that phase information is lost as these pressure waves pass through the inner ear structures of the bat [27], [28], [29]. Indeed, as the bat’s cochlea can be modeled as a set of band pass filters, followed by an envelope detection step, it is apparent that the phase information is effectively lost from the reflected signals. Therefore, the assumption can be made that the inputs into the bat’s auditory system can be adequately modeled by the magnitude of the short-time Fourier transform of the impinging sound pressure waves [30], ignoring the logarithmic spacing of the frequency components in the bat’s auditory system.

In this paper, we aim to build upon this early research and try to lay the foundations for a model for effective 3D scene reconstruction in bats using the required phase-less information. To do this, we let ourselves be inspired by the seminal work achieved in novel view synthesis using deep learning networks. In particular, Neural Radiance Fields (NeRFs), which were first introduced in the seminal paper by Mildenhall et al. at the ECCV conference in 2020 [31]. In that paper, a novel approach for view synthesis is proposed based on building a differentiable visual rendering pipeline, which queries a radiance field represented by a multilayer perceptron (MLP) neural network. The MLP is trained for a specific scene based on images taken from multiple viewpoints, using the differentiable rendering pipeline (DRPL). Novel viewpoints can be generated using the DRPL and the learned radiance function. The original NeRF paper has inspired many researchers and sparked a whole new research field into improving upon the method proposed in the original paper [32], [33], [34]. Furthermore, the usage of NeRFs has been expanded to other application domains such as magnetic resonance imaging [35], [36], ultrasonic medical imaging [37], [38], multimodal acoustic/visual scene representation [39], [40], [41] and acoustic room impulse response prediction [42], with many more other examples to be found.

Based on the success of the underlying concept of NeRFs, more specifically, the differentiable rendering equation combined with a learnable radiance function, we try to explain 3D scene representation by echolocating bats using a NeRF-inspired model. Our model is called a SonoNERF, in which Sono represents ‘sound’ or ‘sonar’, and NERF stands for Neural Reflectivity Field instead of Radiance field, as reflectivity is more appropriate in the context of acoustic echolocation problems. In the remainder of this paper, we will introduce the underlying model of the SonoNERFs and explain how the differentiable rendering pipeline is tailored to the problem of echolocation in bats. Then, we will illustrate the model’s performance on various scenes, and discuss the performance of the model. Finally, we will draw some conclusions and highlight the shortcomings of our model at this point.

## II. Echo formation in echolocating bats

In this section, we will briefly explain the echo formation process in bat echolocation, as this is a requirement to understand the reasoning behind the operating principle of our proposed SonoNERFs. Without loss of generality, we assume that bats emit a broadband signal *s*_*e*_(*t*), which typically is some multi-harmonic chirp [44]. We are aware of the existence of so-called constant-frequency bats. Still, these typically do not perform the gleaning behavior that underlies the SonoNERF principle, so we assume a broadband call. This call is filtered by the external facial features of the bat by a direction-dependent transfer function *h*_*e*_(*ψ, t*), in which *ψ* is a direction vector in 3D, typically represented by the azimuth angle *θ* and elevation angle *φ*. The filtered emitted signal is then reflected by the environment, which we assume to be a collection of point-like reflectors (which, in the Huygens approximation of acoustics, is an acceptable assumption [45]). Each point-reflector filters the impinging signal with a filter *h*_*p*_(*η, t*), characterized b y t he impinging angle *η* (to allow non-isotropic reflection functions to exist in this model). Upon reflection and subsequent reception, the then filtered and reflected signals are filtered by the outer ears of the bats, with a filter *h*_*L*_(*ψ, t*) f or t he l eft ear and a filter *h*_*R*_(*ψ, t*) for the right ear. All these filtered paths are delayed by the round trip range (i.e., from the emitter to the point reflector and from the point reflector to the respective ear).

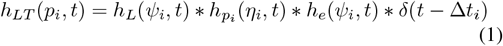

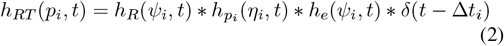

In this equation, the total filters f or a specific po int *p*_*i*_ for the left ear is *h*_*t,L*_(*p*_*i*_, *t*) and for the right ear is *h*_*t,R*_(*p*_*i*_, *t*), and consist of the convolution of a delayed dirac function (*δ*(*t −* Δ*t*_*i*_)), the emission filter *h*_*e*_(*ψ*_*i*_, *t*), the point reflector function *h*_*pi*_(*η*_*i*_, *t*) and the receiver Head Related Transfer Function *h*_*L*_(*ψ*_*i*_, *t*). The received signal for the left and the right ear is the linear sum over all *N* point-like reflectors:

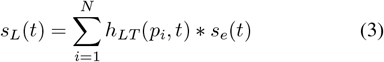

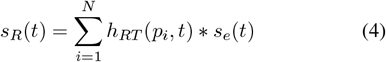

This time-domain representation can be transformed into the Fourier domain, in which the convolutions become multiplications, which facilitates discovering the underlying structure of the physical echo formation process. For this, we first transform the total filters:

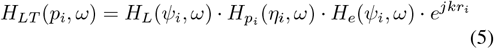

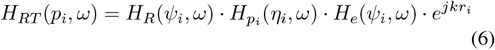

Which we then plug into the equations for the total signals in the left and right ear:

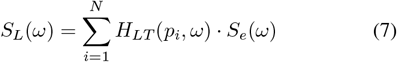

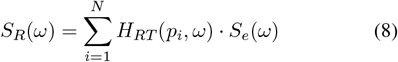

Often, it makes sense to combine the HRTFs of the ears (i.e., *H*_*L*_(*ψ*_*i*_, *ω*)) with the transfer functions of the emitter filter (*H*_*e*_(*ψ*_*i*_, *ω*)) into an object called the ERTF (Echolocation related transfer function, [46], called *E*(*ψ*_*i*_, *ω*):

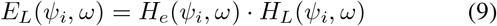

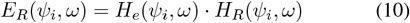

Which in turn reduces the equation for the total signal filter for point *p*_*i*_ to:

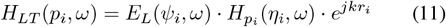

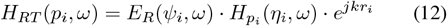

This subtle difference between the HRTF and the ERTF becomes important later in this paper when we describe the differentiable rendering pipeline we implemented for our SonoNERF model. The received signals *s*_*L*_(*t*) and *s*_*R*_(*t*) still contain the phases of the signals. But, as we have argued before, phase information is not considered readily available to bats due to the processing that happens in the bat’s cochlea. Therefore, we approximate the effects of the cochlea by taking the magnitude of the short-term Fourier transform of these received signals:

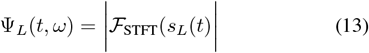

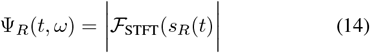

in which ℱ_STFT_ represents the short-term Fourier transform using adequate windowing and overlap values. Subsequently, we concatenate the left and right short-term spectrogram magnitudes into a binaural spectrogram magnitude Ψ_*B*_(*t, ω*):

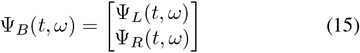

This binaural spectrogram magnitude is then dechirped [47] to remove the time-frequency dependence of the call, which is equivalent to a semi-coherent matched filter (semi-coherent because phase information is not used)[48]:

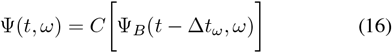

In which the delays Δ*t*_*ω*_ are calculated based on the timefrequency distribution of the emitted signal *s*_*e*_(*t*) [47], [49]. With this, we have arrived at the input data for our SonoNERF: the binaural dechirped magnitude of the short-term Fourier transform Ψ_*D*_(*t, ω*). A nonlinear compression function *C* is applied to map the high dynamic ranges natural to echolocation signals adequately. In the SonoNERF model, a logarithmic compression with linear rescaling was used as a compression function. The matrix Ψ(*t, ω*), together with the pose information of the sensor in the scene, will be the input data for the computations happening inside the SonoNERF model.

## III. Sononerfs

### A. Neural Acoustic Rendering

The NeRF model proposed in [33], solves the task of novel view synthesis. In this task, the model receives a set of observation images *I*_*t*_ and a set of corresponding sensor poses *ζ*_*t*_ consisting of the Cartesian coordinates X, Y, and Z, and the three Euler angles *α, β*, and *γ*. The challenge is synthesizing novel views *I*_*u*_ (or “observations”) for previously unseen poses *ζ*_*u*_. Similarly, the SonoNERF model tries to predict new and unseen dechirped binaural spectrograms Ψ_*u*_(*t, ω*) for previously unseen poses *ζ*_*u*_ based on a training ensemble *E*_*t*_ = [*ζ*_*t*_ Ψ_*t*_]. This prediction step in NeRFs is solved by implementing a differentiable rendering pipeline that uses an underlying radiance field *F*_Θ_ to represent the scene. In many practical applications, this field is represented by a deep multilayer perceptron neural network. Similarly, we will develop a reflectance field *F*_Θ_ for our SonoNERF model, which will, in cooperation with the SonoNERF-DRPL, allow the prediction of novel observations Ψ_*u*_.

In the previous section, we have laid the foundation of the principles governing signal formation in echolocating bats, which we will now adapt to build the SonoNERF-DRPL. The observation Ψ_*u*_ is represented by a 2D matrix of size [2*N*_*f*_ *× N*_*t*_], where *N*_*f*_ is the number of frequency components taken from the STFT (times two because of the binaural concatenation), and *N*_*t*_ is the number of time slices in the STFT, which is the result of the choice of window length and overlap in the STFT computation. In order to calculate the spectrum *ψω* at time *t*, we propose the following differentiable rendering equation:

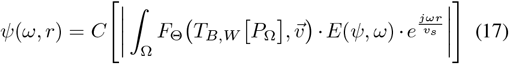

This rendering equation calculates the received spectrum *ψ*_*ω*_ using an integration of hemispheres Ω_*i*_. Figure 1, panel a) shows two of these hemispheres for a certain pose of the bat/sensor. Inside of the integral, the term *E*(*ψ, ω*) denotes the ERTF of the bat, 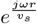 is the Fourier transform of the delay function with *r*_Ω_ the range for the current hemisphere that is being integrated, and *v*_*s*_ is the speed of sound in air. The term *F*_Θ_ (*T*_*B,W*_ [*P*_Ω_], 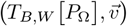) is the neural reflectance field. This function takes as input the position of the voxel for which the reflectivity is to be calculated (*P*_Ω_, converted to world coordinates by transform *T*_*B→W*_), and a direction vector 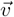, which indicates the direction from which the voxel is being ensonified (which allows non-isotropic reflection surfaces to be modeled with the SonoNERF, similar to the concept of a BRDF in optical rendering [50]). The function *C*() is a nonlinear compression function (as explained in equation 16).

**Fig. 1.**
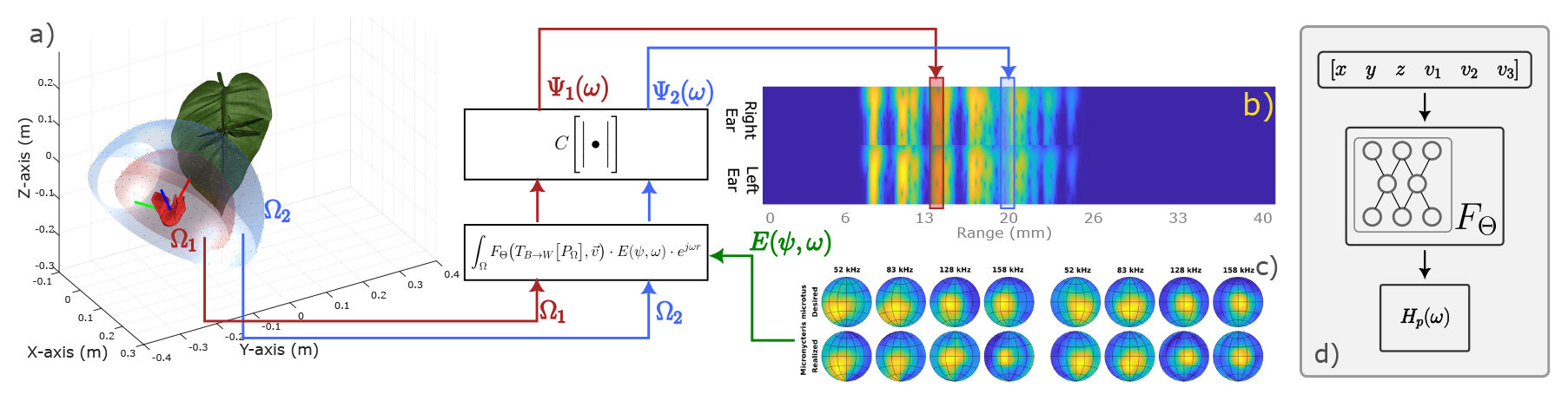
Illustration of the processing flow of the SonoNERF model. Panel a) depicts the bat positioned at a single pose, showcasing two hemispheres Ω used for reflectivity function querying. The reflectivity function is sampled at 600 uniformly distributed points over Ω, which are then used to generate the received spectrum through the rendering equation. This spectrum undergoes nonlinear compression and is placed in the corresponding range slice within the predicted spectrogram (panel b). Panel c) displays the Echolocation-Related Transfer Function (ERTF) for a Micronycteris *microtus* bat call, calculated following the methodology outlined in [43]. Panel d) offers a schematic overview of the Neural Reflectivity field *F*_Θ_, responsible for predicting the reflected spectrum *H*_*p*_(*ω*) based on input position and incidence vector, all represented in world coordinates.

The overall processing flow of this equation can be seen in figure 1. Panel a) shows the bat in a single pose, with two hemispheres Ω. The reflectivity function is queried over a discretized version of this hemisphere (typically, 600 points, distributed uniformly over Ω). The spectrum is calculated through the rendering equation, passed through the nonlinear compression function, and placed at the corresponding range slice in the predicted spectrogram (panel b). Panel c) shows the ERTF for a bat called Micronycteris *microtus* [51], calculated using the method described in [43]. Panel d) shows a schematic representation of the Neural Reflectivity field *F*_Θ_, which predicts the reflected spectrum *H*_*p*_(*ω*) from the input position and incidence vector, all represented in world coordinates.

### B. Training of a SonoNERF

In the previous section, we developed the SonoNERF model, in which a rendering equation was developed to calculate the binaural spectrum received from a scene for a specific range, expressed as *ψ*(*ω, r*). In this rendering equation, a neural reflectivity field represents the scene, called *F*_Θ_, parameterized by a parameter set Θ. In practice, this reflectivity function is implemented using a multi-layer perceptron (MLP), and the parameters Θ are the set of weights and biases for this MLP. In this section, we will detail how this MLP is trained.

Figure 2 shows the overall training process. Data, in the form of a binaural and dechirped spectrogram is recorded from a scene from different poses, as shown in panels a, c, e, and g. The corresponding spectrograms are shown in panels b, d, and f. For a certain pose of the bat and a certain range slice in the spectrogram, we can discretize a hemisphere in front of the bat. This is shown by the blue and red hemispheres (panels a, c, e), which are linked to the corresponding range slices in the spectrogram (panels b, d, f). For each of these hemispheres, we calculate the coordinates of the points on that hemisphere in the world coordinate system, which are then used to query the reflectivity function. This reflectivity function yields a reflectivity value for each of these 3D points and incidence angles, which then is passed through the rendering equation 17 to yield a spectrum *ψ*(*ω*) for a range slice *r*. This predicted spectrum yielded by the rendering function is then compared to the measured spectrum in the spectrogram, from which a loss is calculated (depicted by *ℒ* in figure 2). As both the rendering equation as well as the reflectivity function are fully differentiable, back-propagation can be used to calculate a gradient between the loss *ℒ* and the parameters Θ of the reflectivity function, which allows stochastic gradient descend with back-propagation to be used to optimize these parameters Θ. This approach of training our reflectance function is entirely analogous to the training process of the radiance function described in the original implementation of NeRFs in [31], with the main difference being the type of rendering equation that is used. The versatility of this approach shows the brilliance of the original idea of learnable radiance functions by the authors of [31]. The following section will discuss details on the implementation of the reflectivity function and the training setup.

**Fig. 2.**
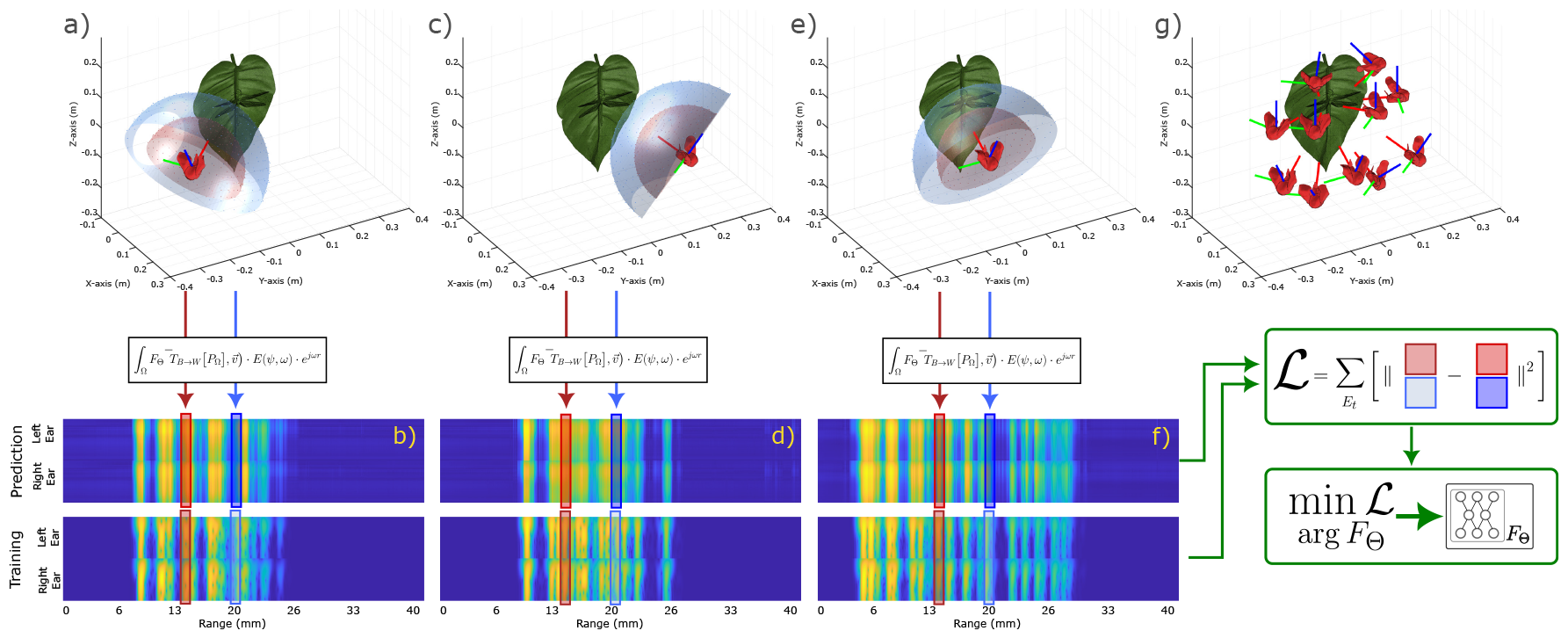
Overview of the SonoNERF Training Process. Data, comprising binaural and dechirped spectrograms, is recorded from various poses within a scene, as depicted in panels a, c, e, and g. The corresponding spectrograms are displayed in panels b, d, and f. Each pose of the bat and range slice in the spectrogram corresponds to a discretized hemisphere in front of the bat, depicted by the blue and red hemispheres in panels a, c, and e, linked to their respective range slices in the spectrogram. Coordinates of points on these hemispheres in the world coordinate system are calculated and used to query the reflectivity function, generating reflectivity values for each 3D point and incidence angle. These values are then processed through rendering equation 17 to produce a spectrum *ψ*(*ω*) for a given range slice *r*. The predicted spectrum is compared to the measured spectrum in the spectrogram, yielding a loss ℒ. This loss ℒ is minimized using stochastic gradient descent by adapting the learnable parameters in function *F*_Θ_.

## IV. Experimental validation

In this section, we will detail the implementation of the SonoNERF setup we developed and show some experimental validation. At this point, we rely purely on simulation to validate the SonoNERF model, as simulation allows us to iterate over ensonification setups rapidly and gives us access to reliable ground truth data. We use SonoTraceLab, a raytracing simulator for simulating acoustic echolocation scenes [52]. The simulator has been extensively validated with real-world experiments, yields reliable data of complex scenes, and allows accurate simulation of biologically relevant acoustic cues. In the remainder of this section, we will provide details on the effective implementation of our SonoNERF model and show its efficacy in multiple experimental scenes. We acknowledge that providing experiments using real-world measured data would be beneficial in illustrating the performance of the SonoNERF approach. However, the authors also believe that frequent scientific communication of relevant bodies of work is crucial to advancing science. Implementing real-world experiments for SonoNERFs is a non-trivial task we will address in future work.

### A.Sononerf Model Implementation

Inside the acoustic rendering equation 17, the neural network *F*_Θ_ represents the neural reflectance field, which encodes the scene being modeled. Figure 3 shows the architecture of this neural network. It consists of a directed acyclic graph and has six input variables: the X, Y, and Z position of the voxel under investigation and the directional vector from where the voxel is being ensonified. The output of the network has 94 values, representing a complex reflectivity function discretized into 47 frequency bins. The first 47 elements of the output of *F* represent the real components of the reflectivity function, and the last 47 elements represent the imaginary components of the reflectivity function. We lift the six input variables inside the network through a Fourier Embedding layer into a higher-dimensional space (6x30 = 180 variables), based on the reasoning provided in [53]. Indeed, embedding low-frequency inputs into a high-frequency representation allows neural networks to learn high-frequency representations better. It should be noted that this step is equivalent to the approach taken in the original NeRF implementation [33].

**Fig. 3.**
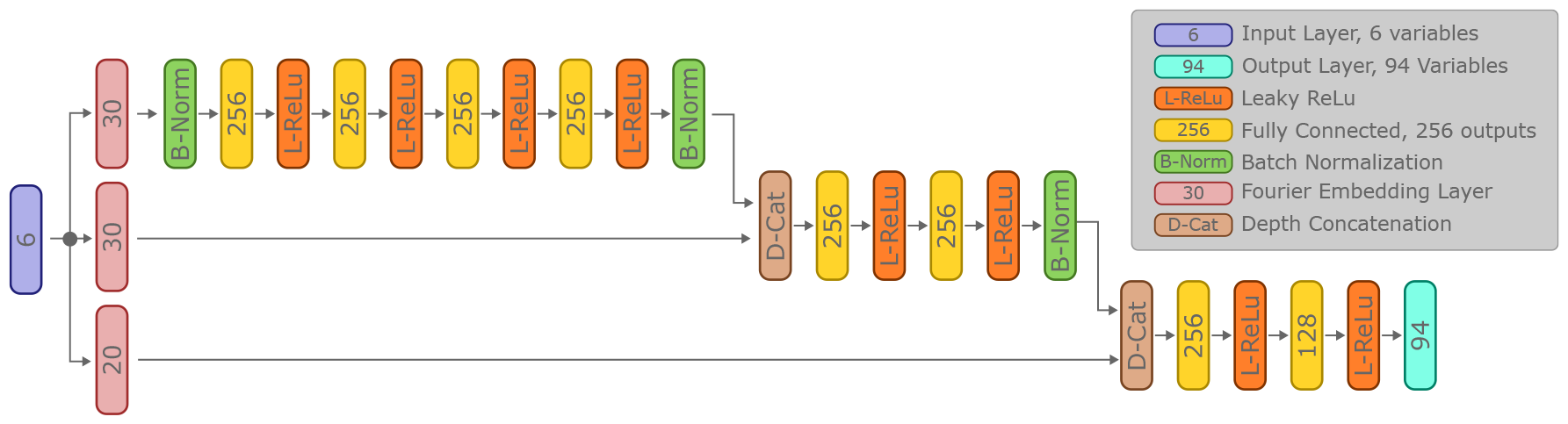
Overview of the SonoNERF reflectivity function *F*_Θ_, which encodes a scene’s acoustic properties within a neural reflectance field framework. The network takes six input variables: the *X, Y*, and *Z* coordinates of a voxel, along with a directional vector indicating the angle of ensonification. These inputs undergo a Fourier Embedding process, expanding them into a higher-dimensional space (180 variables) to capture high-frequency details more effectively. The network architecture features multiple fully connected layers with Leaky ReLU activation functions and incorporates skip connections for enhanced gradient flow during training. The network output consists of 94 values, which describe a complex reflectivity function across 47 frequency bins, with the first 47 values representing the real components and the last 47 the imaginary components of the reflectivity function.

After the Fourier embedding step, the network consists of several fully connected layers, combined with Leaky ReLu activation functions as non-linearity [54]. Skip connections are added to the network to allow for more efficient gradient flow, allowing faster learning during training. Skip connections are concatenated through depth concatenation, increasing the size of the inputs of the subsequent layer following the depth concatenation.

We implemented the SonoNERF model, including the rendering equation and the neural reflectance function, in Matlab using object-oriented programming methods and used the Deep Learning toolbox [55] to perform automatic differentiation on the complete rendering equation. This allows rapid iteration over multiple versions of the rendering equation without manually calculating the gradients of the rendering equation. We optimized the performance of the rendering equations by using the GPUArray objects in Matlab, which allow rapid evaluation of matrix and vector operations on the GPU of the system. This allows data to stay in GPU memory during the evaluation of the rendering equation, which yields a significant speed-up compared to a naive CPU implementation (in our tests, speedups of around 100x were achieved).

Training of the SonoNERF models was performed using Adam [56], with a batch size of 512, a learning rate of 0.01, and a learning rate drop factor of 0.97. We trained the networks for 100 epochs using a single NVidia RTX4090, which took around 7 hours to complete. The network described in figure 3 has around 500.000 learnable parameters.

### B. From Spectrograms to 3D Scene Description

The SonoNERF model is a method to predict the magnitude of spectrograms that would be observed for previously unseen poses using a model that is trained using measured magnitude spectrograms observed from a set of training poses. Once trained, the SonoNERF model can predict a new spectrogram for an arbitrary pose and nothing more. Indeed, from the point of view of the training algorithm, the concept of a 3D scene description is not made explicit during the training process. However, it is possible to query the trained reflectance function to obtain a 3D voxel description of the environment [33]. To achieve this reconstruction of the 3D scene geometry, we discretize the volume of interest in a cubic voxel with a side of 0.5mm and use 100 ensonification directions distributed uniformly over the sphere for each voxel. We then query the trained reflectance *F*_*T*_ *heta* for each of these voxel and ensonification directions. The result is a matrix *R* with dimensions:

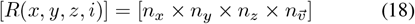

with *n*_*x*_, *n*_*y*_, and *n*_*z*_ the number of voxels in the X, Y, and Z dimensions, respectively, and 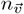 being the number of sampled ensonification d irections (100 i n o ur e xamples). N ext, we integrate this reflectivity matrix over the direction dimension:

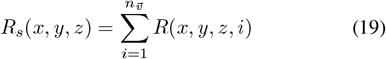

Next, using a fixed threshold, we calculate the isosurface over this function *R*_*s*_(*x, y, z*). This yields a surface *V*_*r*_, which can then be visualized. It should be noted that various other visualization methods could be used, such as maximum intensity projection [57] or a plethora of other techniques [58].

However, for the scoping of this paper we opted for applying the isosurface method.

### C. SonoNERF trained on a simple scene

In order to validate the proposed SonoNERF model, we experimented using a simple scene. The model used in this experiment is shown in figure 4, panel a). This panel shows three spheres with a diameter of 3cm, arranged in an Lshaped configuration. The figure also shows three poses (which were not part of the training set), ensonifying the scene. We used 100 observations spaced uniformly around the scene and calculated the received spectrograms using our SonoTraceLab simulator [52]. This simulator uses a ray-acoustics based approach and is ideally suited to simulate complex echolocation scenes, and has been validated using real-world experiments. The simulated spectrograms can be seen in the top row of panels b, d, and f. One can see various direct and multi-path reflections occurring later in time, which is expected based on the scene geometry. SonoTraceLab is capable of simulating complex multi-path reflections occurring in challenging scenes. After training, the SonoNERF model is capable of reconstructing the spectrograms for previously unseen poses, shown in panels b, d, and f, bottom row. Both the location as well as spectral content of these reconstructed spectrograms correspond to the ground truth that was simulated. Panel c) shows a magnified view of the scene, and panel e) shows the isosurface *V*_*r*_ reconstructed from the queried reflectance function *F*_Θ_ accumulated into *R*. Panel g) shows a maximum intensity projection of the same reflectivity matrix *R*. The main shape and location of the individual spheres are reconstructed, while the size of the spheres appears to be inflated. This inflation most likely happens due to the absence of phase information in the input data, which diminishes range resolution because only semi-coherent matched filters can be used [30].

**Fig. 4.**
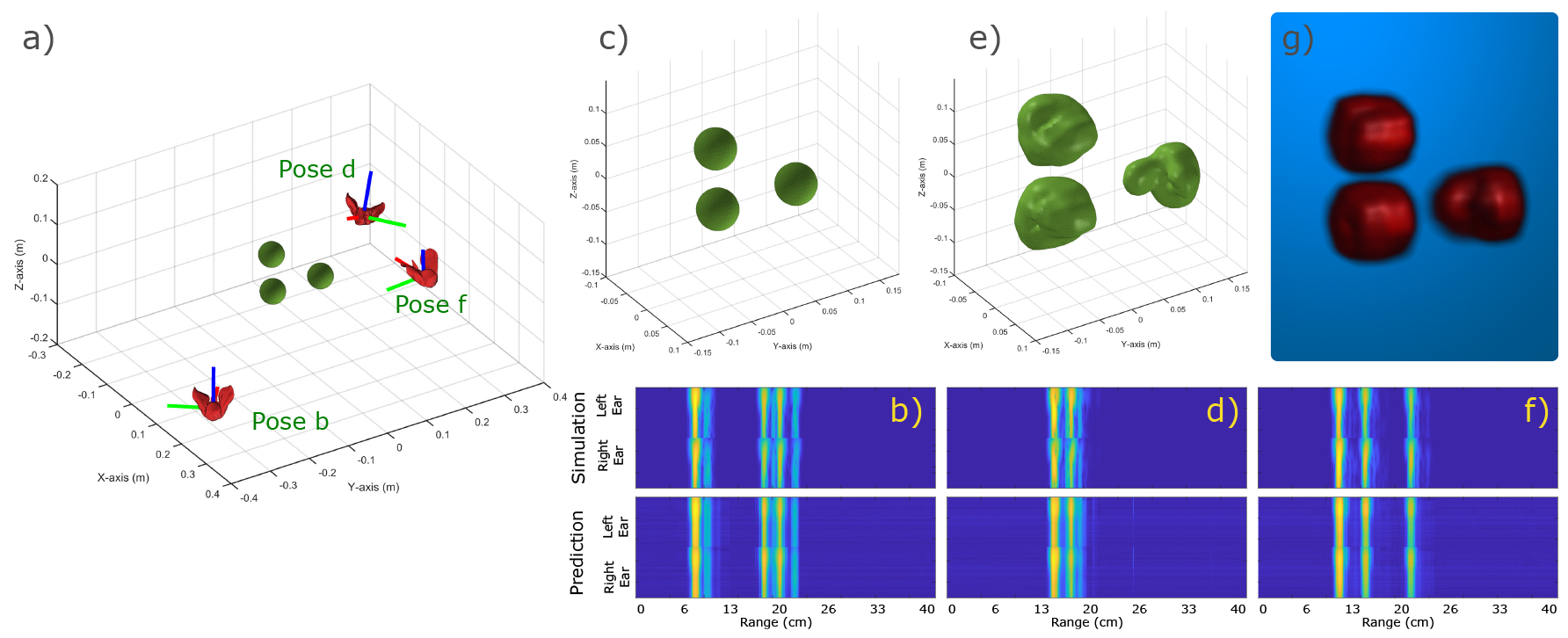
Overview of the result of a trained SonoNERF model on a simple scene. Panel a) shows the scene, which consists of three spheres in an L-shaped configuration. Three poses (b, d, and f) are shown from which the scene is ensonified. The corresponding received spectrograms are shown in panels b, d, and f, top panels (called “simulation”). We trained the described SonoNERF model using 100 observations. The resulting predicted spectrograms for poses b, d, and f (not part of the training set) are shown in the top row of panels b, d, and f. Panel c) shows a more magnified scene plot, and panel e) shows the reconstructed isosurface *V*_*r*_ . Panel g) shows a maximum intensity projection of the same reconstruction.

### D. SonoNERF trained on a complex scene

After our initial validation of the SonoNERF model, we constructed a more complicated scene. This scene can be seen in figure 5. This scene consists of 19 spheres arranged to form the letters “UA”, corresponding to the name of our institute, University of Antwerp. Similar to the previous subsection, we show the simulated and reconstructed spectrograms. It becomes clear that the time separation between the individual reflections in the spectrograms is less clear, causing the echoes to overlap more. This is also reflected in the reconstructed isosurface, shown in panel e). The individual spheres are no longer separated but form a continuous surface. However, despite these shortcomings, the letters UA are recognizable, with the important features of the letters being conserved (such as the gap between the vertical parts of the U or the hole inside of the A).

**Fig. 5.**
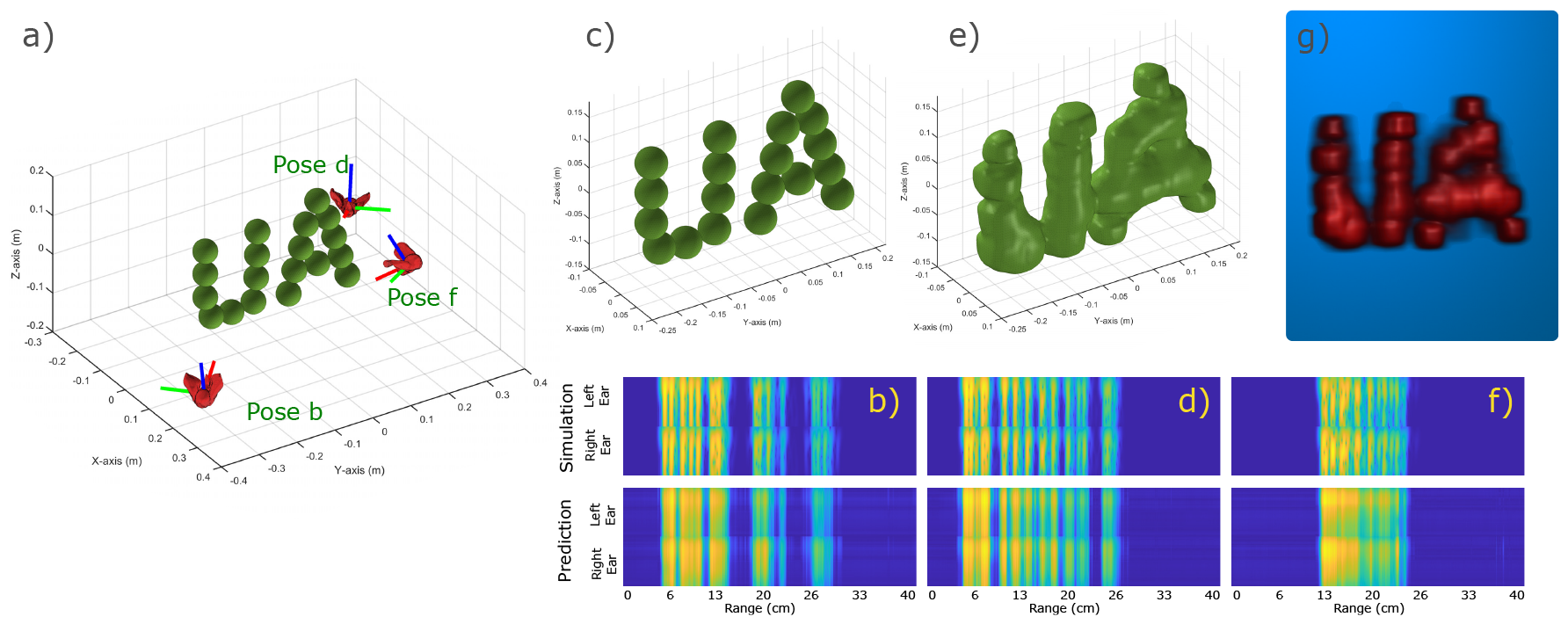
Overview of the result of a trained SonoNERF model on a more complex scene. Panel a) shows the scene, which consists of 19 spheres arranged to form the letters “UA”, for the University of Antwerp. Three poses (b, d, and f) are shown from which the scene is ensonified. The corresponding received spectrograms are shown in panels b, d, and f, top panels (called “simulation”). We trained the described SonoNERF model using 100 observations. The resulting predicted spectrograms for the poses b, d, and f (not part of the training set) are shown in the top row of panels b, d, and f. Panel c) shows a more magnified scene plot, and panel e) shows the reconstructed isosurface *V*_*r*_ . Panel g) shows a maximum intensity projection of the same reconstruction.

### E. SonoNERF trained on a biologically relevant scene

As a final demonstration of the capabilities of our SonoNERF model, we constructed a scene consisting of a leaf on which we perched a dragonfly, which is a scenario that is performed by hunting Micronycteris *microtus* bats in the neotropics [59]. We performed the same simulation and SonoNERF training as in the two previous scenarios, and show the results in figure 6. The SonoNERF is capable in reconstructing the spectrogram representations (panels b, d, f), as well as the overall leaf geometry (panels e and g). To further investigate this scenario, we modelled the leaf with and without a dragonfly present on the leaf surface. The result of this reconstruction can be seen in figure 7. Panel a) shows the leaf without dragonfly, and panel c) shows the leaf with dragonfly. Panels b) and d) show the isosurface reconstruction of these two scenarios. Here, a clear bulge can be seen in the reconstruction of the leaf with dragonfly (panel d), which is absent in the reconstruction of the leaf without dragonfly (panel b). Panel e) shows the maximum intensity projection of energy matrix *R*_*s*_ (eq. 19) into the direction of the camera viewpoint, which have been normalized to the maximum of both *R*_*s*_ for the two reconstructions. It becomes apparent that the overall reflection strength of the leaf with dragonfly is much larger compared to the leaf without dragonfly. The top-view of this representation (panel f) further details this. Finally, panel g) shows the difference in energy between the reflected energy matrix of the leaf with and without dragonfly. Here, a strong energy peak can be seen in the middle of the leaf surface. To further illustrate this energy difference, we plotted the energy difference using maximum intensity projection on top of the STL model of the leaf with dragonfly. This can be seen in figure 8, panel c). In this overlay, a strong peaks can be observed around the location of the dragonfly on the leaf.

**Fig. 6.**
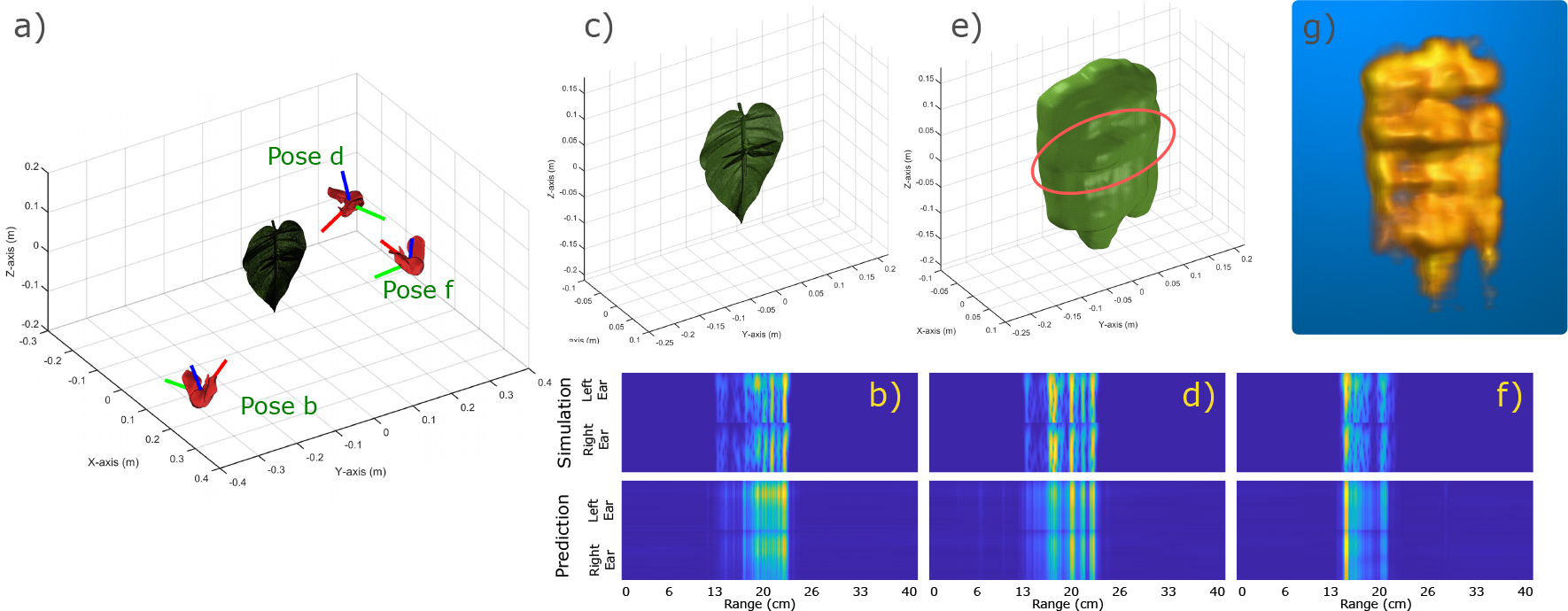
Overview of the result of a trained SonoNERF model on a biologically relevant scene of an insect perched on a leaf. Panel a) shows the scene, a leaf with a dragonfly, similar to the approaches of Micronycteris *microtus* like in [59]. Three poses (b, d, and f) are shown from which the scene is ensonified. The corresponding received spectrograms are shown in panels b, d, and f, top panels (called “simulation”). We trained the described SonoNERF model using 100 observations. The resulting predicted spectrograms for the poses b, d, and f (not part of the training set) are shown in the top row of panels b, d, and f. Panel c) shows a more magnified scene plot, and panel e) shows the reconstructed isosurface *V*_*r*_ . Panel g) shows a maximum intensity projection of the same reconstruction. Panel e) shows a thickening in the volume mesh on the location of the dragonfly, hinting for detailed scene information to be present in the reconstructed mesh.

**Fig. 7.**
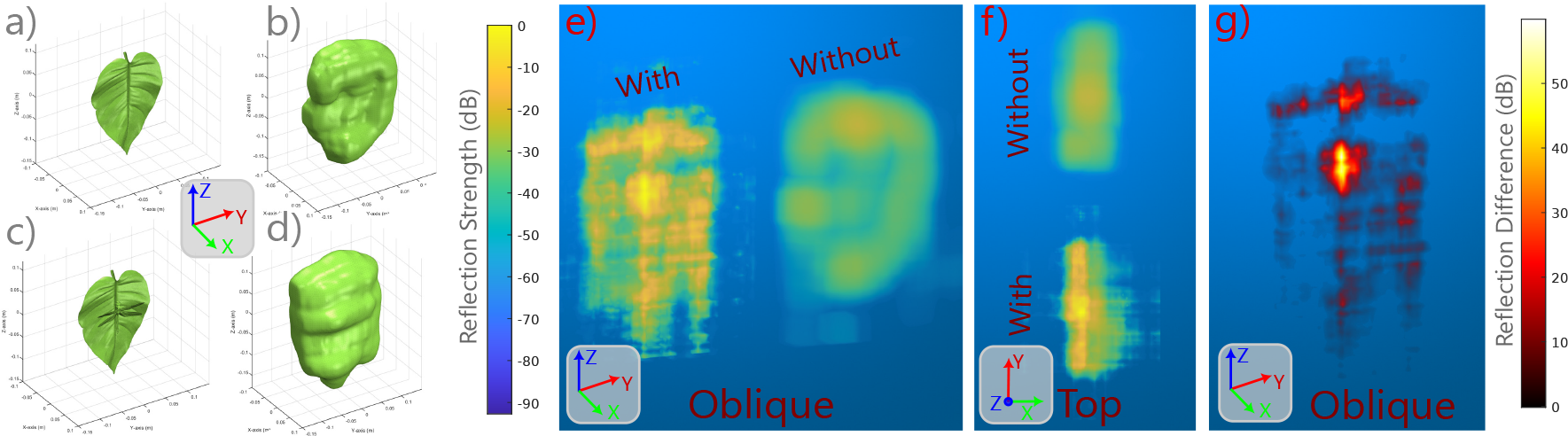
Overview of the SonoNERF volume reconstructions for a leaf without an insect (panels a and b) and for a leaf with an insect (panels c and d). A significant bulge in the mesh surface can be observed on the location of the dragonfly in panel d). Panel e) shows the reflectivity function through maximum intensity projection (MIP) into the camera, normalized to the strongest reflection across the two instances. Panel f) shows the top-view of the same MIP visualizatio. Finally, panel g) shows the difference in reflectivity function for a leaf with and without dragonfly.

**Fig. 8.**
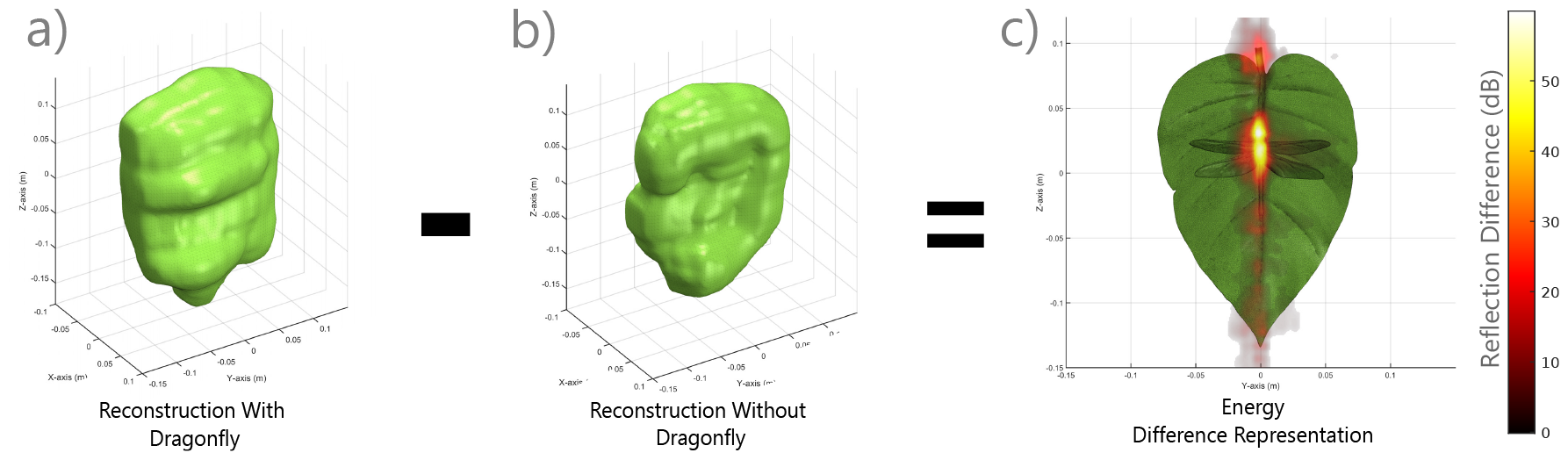
Detail of the difference in reflectivity function between a leaf with dragonfly and without dragonfly. The largest differences can be observed around the location of the dragonfly where the difference function peaks strongly. This could be used as a cue for prey localization on the leaf.

These results illustrate a potential biological implication of our SonoNERF model. As shown, the SonoNERF reconstruction allows for the reconstruction of biologically relevant information from the scene: it allows the bat to localize the dragonfly on the leaf, and to distinguish between a filled and an empty leaf. While the model proposed in [59] for leaf occupancy state discrimination has not lost it’s validity, the model therein proposed does not explain how the bat might be able to localize the prey item on the leaf. While me make no claims on which model is implemented in the bats brain, we do think that With our proposed SonoNERF model, we have shown that prey localization against background clutter is enabled by such a data-aggregation task. More specifically, through learning a prediction task of novel spectrograms from unseen poses, the surface reconstruction, and therefore the prey localization, emerges as a byproduct of the computational graph that is used to solve the prediction task.

## V. Discussion and conclusion

In this paper, we proposed SonoNERFs, a novel model for 3D scene reconstruction in a biologically plausible manner. We use the concept of neural radiance fields to solve the problem of predicting echo spectrograms that would be obtained from a scene when ensonified from previously unseen poses, without access to the phase information of the received echoes. As explained, phase coherence and phase reception are unlikely to exist in echolocating bats because of how vocalization and reception (most notably the cochlea) behave in real animals. After we provided a solution for the spectrogram prediction problem, we detailed how the learned reflectivity model can be used to perform 3D scene reconstruction of complex shapes, which we demonstrated using three scenes of varying complexity.

One could argue that the fact that 3D scene reconstruction becomes possible when combining measurements from different poses is not that surprising. Indeed, computed tomography techniques exist, and are already applied to echolocation scenarios. However, these systems utilize phase-coherent measurements and need the phase information of the echoes to work well. What, in the opinion of the authors, is not so trivial is the fact that 3D scene reconstruction emerges when solving a completely different task, namely predicting sensor data for a novel scene. While it has been observed that bats can predict aspects of the scene by accumulating sensor data [60], to the best of our knowledge, no concrete model on how this prediction might operate has been proposed in previous works. Other technical systems have been proposed to produce 3D scene reconstruction and semantic interpretation, [61], [62], but these proposed techniques utilize a teaching modality like LIDARs or cameras to perform a form of modality translation. Our SonoNERF model relies solely on acoustic data without the need for an additional supervision modality. Furthermore, reference [62] does not use an acoustic sensing modality, causing the title to be somewhat misleading. Our approach follows the approach called ‘self-supervised learning’ which has received much attention in the recent years [63], [64]. Through self-supervision, computational graphs can learn how to predict their own inputs, and, based on the structure of the implemented computational graph, can lead to emergent insights into the underlying sensor data (such as demonstrated in our SonoNERF model).

In addition to the capability of SonoNERF to predict spectrograms and to perform surface reconstruction, we also showed how the SonoNERF model could be used to solve a relevant task for a biological echolocator, namely, finding motionless prey items against strong clutter backgrounds. While we postulated a model in [59] on how the bat Micronycteris *microtus* might be able to distinguish between an empty or a full leaf, no concrete postulate was made on how localization might take place. In the previous section, we demonstrated how a SonoNERF model might be used to perform exactly this task, through the examination of the reconstructed reflectivity function. It should be noted that while our proposed SonoNERF model might be one solution to how scene reconstruction takes place, we nowhere claim that this is the exact model that is implemented in the brain of a bat. More specifically, what we do claim is that the SonoNERF model is a first-order hypothesis to how bats might be able to solve this problem, similarly like we proposed our BatSLAM algorithm as a first-order model on how bats might perform large-scale localization [65]. In-depth biological behaviour experiments need to be performed which use testable hypotheses generated through our SonoNERF model to gain insight into the true underlying behavioral mechanisms.

We acknowledge the lack of real-world measurements in this paper. However, as we argued before, using our validated SonoTraceLab simulator is a strong alternative approach to producing experimental data, as this simulator has been demonstrated to be able to produce biologically relevant acoustic data from complex scenes without the arising complexities of performing real-world data. We also acknowledge that real-world experiments are relevant, which is why these experiments are part of our short-term future work. Using SonoTraceLab to provide us with data allows us to rapidly iterate over experiments, and allows complex visualizations such as the one in figure 8, where we overlay the reconstructed reflectivity function over the base model, which then allows the discovery of important features such as the difference of a leaf with and without insect.

Next to performing real-world experimentation, several improvements can be proposed to the proposed model. For example, at this point our SonoNERF model is trained from a randomly initialized reflectivity model each time. One could argue that, over time, bats learn priors on how scenes might be encoded, which could then be reflected into prior models or transfer learned models later. Furthermore, the acoustic rendering equation 17 has no concept of multi-path signal propagation. One natural extension would be to expand this rendering equation to encompass multi-path reflections. However, this augmented rendering equation should still be fully differentiable, which is currently unknown to the authors whether this will be the case.

## Notes

### Competing Interest Statement

The authors have declared no competing interest.

## References

[1] G. P. Bell, “Behavioral and ecological aspects of gleaning by a desert insectivorous bat Antrozous pallidus (Chiroptera: Vespertilionidae),” Behavioral ecology and sociobiology, vol. 10, pp. 217–223, 1982.

[2] A. C. Entwistle, P. A. Racey, and J. R. Speakman, “Habitat exploitation by a gleaning bat, Plecotus auritus,” Philosophical Transactions of the Royal Society of London. Series B: Biological Sciences, vol. 351, no. 1342, pp. 921–931, 1996.

[3] I. Geipel, K. Jung, and E. K. Kalko, “Perception of silent and motionless prey on vegetation by echolocation in the gleaning bat Micronycteris microtis,” Proceedings of the Royal Society B: Biological Sciences, vol. 280, no. 1754, p. 20122830, 2013.

[4] K. A. Razak, “Adaptations for substrate gleaning in bats: The pallid bat as a case study,” Brain, behavior and evolution, vol. 91, no. 2, pp. 97–108, 2018.

[5] S. Stoffberg and D. S. Jacobs, “The influence of wing morphology and echolocation on the gleaning ability of the insectivorous bat Myotis tricolor,” Canadian journal of zoology, vol. 82, no. 12, pp. 1854–1863, 2004.

[6] S. Swift and P. Racey, “Gleaning as a foraging strategy in Natterer’s bat Myotis nattereri,” Behavioral Ecology and Sociobiology, vol. 52, pp. 408–416, 2002.

[7] I. Geipel, J. Steckel, M. Tschapka, D. Vanderelst, H.-U. Schnitzler, E. K. Kalko, H. Peremans, and R. Simon, “Bats actively use leaves as specular reflectors to detect acoustically camouflaged prey,” Current biology, vol. 29, no. 16, pp. 2731–2736, 2019.

[8] E. Verreycken, R. Simon, B. Quirk-Royal, W. Daems, J. Barber, and J. Steckel, “Bio-acoustic tracking and localization using heterogeneous, scalable microphone arrays,” Communications Biology, vol. 4, no. 1, p. 11 p., 2021.

[9] R. Arlettaz, G. Jones, and P. A. Racey, “Effect of acoustic clutter on prey detection by bats,” Nature, vol. 414, no. 6865, pp. 742–745, 2001.

[10] B. M. Siemers, E. Baur, and H.-U. Schnitzler, “Acoustic mirror effect increases prey detection distance in trawling bats,” Naturwissenschaften, vol. 92, pp. 272–276, 2005.

[11] S. Zsebok, F. Kroll, M. Heinrich, D. Genzel, B. M. Siemers, and L. Wiegrebe, “Trawling bats exploit an echo-acoustic ground effect,” Frontiers in Physiology, vol. 4, p. 65, 2013.

[12] T. U. Grafe, C. R. Schöner, G. Kerth, A. Junaidi, and M. G. Schöner, “A novel resource–service mutualism between bats and pitcher plants,” Biology Letters, vol. 7, no. 3, pp. 436–439, 2011.

[13] M. G. Schöner, C. R. Schöner, R. Simon, T. U. Grafe, S. J. Puechmaille, L. L. Ji, and G. Kerth, “Bats are acoustically attracted to mutualistic carnivorous plants,” Current Biology, vol. 25, no. 14, pp. 1911–1916, 2015.

[14] R. Simon, K. Bakunowski, A. E. Reyes-Vasques, M. Tschapka, M. Knoernschild, J. Steckel, and D. Stowell, “Acoustic traits of bat-pollinated flowers compared to flowers of other pollination syndromes and their echo-based classification using convolutional neural networks,” PLoS computational biology, vol. 17, no. 12, p. 20 p., 2021.

[15] R. Simon, F. Matt, V. Santillan, M. Tschapka, M. Tuttle, and W. Halfwerk, “An ultrasound-absorbing inflorescence zone enhances echo-acoustic contrast of bat-pollinated cactus flowers,” Journal of Experimental Biology, vol. 226, no. 5, p. jeb245263, 2023.

[16] R. Simon, S. Rupitsch, M. Baumann, H. Wu, H. Peremans, and J. Steckel, “Bioinspired sonar reflectors as guiding beacons for autonomous navigation,” Proceedings of the National Academy of Sciences of the United States of America, vol. 117, no. 3, pp. 1367–1374, 2020.

[17] M. de Backer, W. Jansen, D. Laurijssen, R. Simon, W. Daems, and J. Steckel, “Detecting and classifying bio-inspired artificial landmarks using in-air 3D sonar,” in 2023 IEEE SENSORS, 29 October - 01 November, 2023, Vienna, Austria. IEEE, 2023, pp. 1–4.

[18] M. Denny, “The physics of bat echolocation: Signal processing techniques,” American Journal of Physics, vol. 72, no. 12, pp. 1465–1477, 2004.

[19] R. A. Altes, “Sonar for generalized target description and its similarity to animal echolocation systems,” The Journal of the Acoustical Society of America, vol. 59, no. 1, pp. 97–105, 1976.

[20] P. A. Saillant, J. A. Simmons, S. P. Dear, and T. A. McMullen, “A computational model of echo processing and acoustic imaging in frequency-modulated echolocating bats: The spectrogram correlation and transformation receiver,” The Journal of the Acoustical Society of America, vol. 94, no. 5, pp. 2691–2712, 1993.

[21] J. A. Simmons and R. A. Stein, “Acoustic imaging in bat sonar: Echolocation signals and the evolution of echolocation,” Journal of comparative physiology, vol. 135, pp. 61–84, 1980.

[22] J. A. Simmons, C. F. Moss, and M. Ferragamo, “Convergence of temporal and spectral information into acoustic images of complex sonar targets perceived by the echolocating bat, Eptesicus fuscus,” Journal of Comparative Physiology A, vol. 166, pp. 449–470, 1990.

[23] J. A. Simmons, “A view of the world through the bat’s ear: The formation of acoustic images in echolocation,” Cognition, vol. 33, no. 1-2, pp. 155–199, 1989.

[24] A. Balleri, H. D. Griffiths, K. Woodbridge, C. J. Baker, and M. W. Holderied, “Bat-inspired ultrasound tomography in air,” in 2010 IEEE Radar Conference. IEEE, 2010, pp. 44–47.

[25] E. L. Clare and M. W. Holderied, “Acoustic shadows help gleaning bats find prey, but may be defeated by prey acoustic camouflage on rough surfaces,” Elife, vol. 4, p. e07404, 2015.

[26] T. R. Neil, Z. Shen, D. Robert, B. W. Drinkwater, and M. W. Holderied, “Moth wings are acoustic metamaterials,” Proceedings of the National Academy of Sciences, vol. 117, no. 49, pp. 31 134–31 141, 2020.

[27] A. Chitradurga Achutha, H. Peremans, U. Firzlaff, and D. Vanderelst, “Efficient encoding of spectrotemporal information for bat echolocation,” PLOS Computational Biology, vol. 17, no. 6, p. e1009052, 2021.

[28] S. Y. Kim, R. Allen, and D. Rowan, “The simulation of bat-oriented auditory processing using the experimental data of echolocating signals,” Journal of the Acoustical Society of America, vol. 123, no. 5, p. 3621, 2008.

[29] M. Kössl and M. Vater, “The cochlear frequency map of the mustache bat, Pteronotus parnellii,” Journal of Comparative Physiology A, vol. 157, pp. 687–697, 1985.

[30] H. Peremans and J. Hallam, “The spectrogram correlation and transformation receiver, revisited,” The Journal of the Acoustical Society of America, vol. 104, no. 2, pp. 1101–1110, 1998.

[31] B. Mildenhall, P. P. Srinivasan, M. Tancik, J. T. Barron, R. Ramamoorthi, and R. Ng, “Nerf: Representing scenes as neural radiance fields for view synthesis,” Communications of the ACM, vol. 65, no. 1, pp. 99–106, 2021.

[32] K. Zhang, G. Riegler, N. Snavely, and V. Koltun, “Nerf++: Analyzing and improving neural radiance fields,” arXiv preprint 2010.07492, 2020.

[33] K. Gao, Y. Gao, H. He, D. Lu, L. Xu, and J. Li, “Nerf: Neural radiance field in 3d vision, a comprehensive review,” arXiv preprint 2210.00379, 2022.

[34] F. Zhu, S. Guo, L. Song, K. Xu, and J. Hu, “Deep review and analysis of recent nerfs,” APSIPA Transactions on Signal and Information Processing, vol. 12, no. 1, 2023.

[35] K. Iddrisu, S. Malec, and A. Crimi, “3D reconstructions of brain from MRI scans using neural radiance fields,” in International Conference on Artificial Intelligence and Soft Computing. Springer, 2023, pp. 207–218.

[36] T. J. Jang and C. M. Hyun, “NeRF Solves Undersampled MRI Reconstruction,” arXiv preprint 2402.13226, 2024.

[37] M. Wysocki, M. F. Azampour, C. Eilers, B. Busam, M. Salehi, and N. Navab, “Ultra-nerf: Neural radiance fields for ultrasound imaging,” in Medical Imaging with Deep Learning. PMLR, 2024, pp. 382–401.

[38] Y. Zou, Y. Lin, and Q. Zhu, “PA-NeRF, a neural radiance field model for 3D photoacoustic tomography reconstruction from limited Bscan data,” Biomedical Optics Express, vol. 15, no. 3, pp. 1651–1667, 2024.

[39] C. Chen, A. Richard, R. Shapovalov, V. K. Ithapu, N. Neverova, K. Grauman, and A. Vedaldi, “Novel-view acoustic synthesis,” in Proceedings of the IEEE/CVF Conference on Computer Vision and Pattern Recognition, 2023, pp. 6409–6419.

[40] Z. Chen, I. D. Gebru, C. Richardt, A. Kumar, W. Laney, A. Owens, and A. Richard, “Real Acoustic Fields: An Audio-Visual Room Acoustics Dataset and Benchmark,” arXiv preprint 2403.18821, 2024.

[41] Y. Guo, K. Chen, S. Liang, Y.-J. Liu, H. Bao, and J. Zhang, “Adnerf: Audio driven neural radiance fields for talking head synthesis,” in Proceedings of the IEEE/CVF International Conference on Computer Vision, 2021, pp. 5784–5794.

[42] A. Luo, Y. Du, M. Tarr, J. Tenenbaum, A. Torralba, and C. Gan, “Learning neural acoustic fields,” Advances in Neural Information Processing Systems, vol. 35, pp. 3165–3177, 2022.

[43] F. De Mey, J. Reijniers, H. Peremans, M. Otani, and U. Firzlaff, “Simulated head related transfer function of the phyllostomid bat Phyllostomus discolor,” The Journal of the Acoustical Society of America, vol. 124, no. 4, pp. 2123–2132, 2008.

[44] G. Jones and M. W. Holderied, “Bat echolocation calls: Adaptation and convergent evolution,” Proceedings of the Royal Society B: Biological Sciences, vol. 274, no. 1612, pp. 905–912, 2007.

[45] A. D. Pierce, Acoustics: An Introduction to Its Physical Principles and Applications. Springer, 2019.

[46] J. Steckel and H. Peremans, “A novel biomimetic sonarhead using beamforming technology to mimic bat echolocation,” IEEE transactions on ultrasonics, ferroelectrics and frequency control, vol. 59, no. 7, pp. 1369–1377, 2012.

[47] J. Wang, D. Cai, and Y. Wen, “Comparison of matched filter and dechirp processing used in linear frequency modulation,” in 2011 IEEE 2nd International Conference on Computing, Control and Industrial Engineering, vol. 2. IEEE, 2011, pp. 70–73.

[48] L. Wiegrebe, “An autocorrelation model of bat sonar,” Biological cybernetics, vol. 98, pp. 587–595, 2008.

[49] J. Steckel, D. Vanderelst, and H. Peremans, “BatSLAM : Combining biomimetic sonar with a hippocampal model,” in Proceedings of the Robotica Conference, Guimaraes, Portugal, 2012, pp. –.

[50] W. Matusik, H. Pfister, M. Brand, and L. McMillan, “Efficient isotropic BRDF measurement,” 2003.

[51] D. Vanderelst, F. De Mey, H. Peremans, I. Geipel, E. Kalko, and U. Firzlaff, “What noseleaves do for FM bats depends on their degree of sensorial specialization,” PloS one, vol. 5, no. 8, p. e11893, 2010.

[52] W. Jansen and J. Steckel, “SonoTraceLab-A Raytracing-Based Acoustic Modelling System for Simulating Echolocation Behavior of Bats,” arXiv preprint 2403.06847, 2024.

[53] M. Tancik, P. Srinivasan, B. Mildenhall, S. Fridovich-Keil, N. Raghavan, U. Singhal, R. Ramamoorthi, J. Barron, and R. Ng, “Fourier features let networks learn high frequency functions in low dimensional domains,” Advances in neural information processing systems, vol. 33, pp. 7537–7547, 2020.

[54] C. Banerjee, T. Mukherjee, and E. Pasiliao Jr, “An empirical study on generalizations of the ReLU activation function,” in Proceedings of the 2019 ACM Southeast Conference, 2019, pp. 164–167.

[55] “Deep Learning Toolbox,” https://nl.mathworks.com/products/deep-learning.html.

[56] D. P. Kingma and J. Ba, “Adam: A method for stochastic optimization,” arXiv preprint 1412.6980, 2014.

[57] S. Napel, M. P. Marks, G. D. Rubin, M. D. Dake, C. H. McDonnell, S. M. Song, D. R. Enzmann, and R. B. Jeffrey Jr, “CT angiography with spiral CT and maximum intensity projection.” Radiology, vol. 185, no. 2, pp. 607–610, 1992.

[58] A. E. Kaufman and K. Mueller, “Overview of volume rendering.” The visualization handbook, vol. 7, pp. 127–174, 2005.

[59] I. Geipel, J. Steckel, M. Tschapka, D. Vanderelst, H.-U. Schnitzler, E. Kalko, H. Peremans, and R. Simon, “Bats actively use leaves as specular reflectors to detect acoustically camouflaged prey,” Current Biology, vol. 29, no. 16, 2019.

[60] A. Salles, C. A. Diebold, and C. F. Moss, “Echolocating bats accumulate information from acoustic snapshots to predict auditory object motion,” Proceedings of the National Academy of Sciences, vol. 117, no. 46, pp. 29 229–29 238, 2020.

[61] J. H. Christensen, S. Hornauer, and X. Y. Stella, “Batvision: Learning to see 3d spatial layout with two ears,” in 2020 IEEE International Conference on Robotics and Automation (ICRA). IEEE, 2020, pp. 1581–1587.

[62] C. Zhang, Z. Yang, B. Xue, H. Zhuo, L. Liao, X. Yang, and Z. Zhu, “Perceiving like a Bat: Hierarchical 3D Geometric–Semantic Scene Understanding Inspired by a Biomimetic Mechanism,” Biomimetics, vol. 8, no. 5, p. 436, 2023.

[63] D. Hendrycks, M. Mazeika, S. Kadavath, and D. Song, “Using self-supervised learning can improve model robustness and uncertainty,” Advances in neural information processing systems, vol. 32, 2019.

[64] X. Liu, F. Zhang, Z. Hou, L. Mian, Z. Wang, J. Zhang, and J. Tang, “Self-supervised learning: Generative or contrastive,” IEEE transactions on knowledge and data engineering, vol. 35, no. 1, pp. 857–876, 2021.

[65] J. Steckel and H. Peremans, “BatSLAM: Simultaneous Localization and Mapping Using Biomimetic Sonar,” PLoS ONE, vol. 8, no. 1, Jan. 2013.

